# Comparison of two descriptions of heterogeneity used in modelling plants dynamics

**DOI:** 10.1101/075812

**Authors:** Michel Droz, Andrzej Pękalski

## Abstract

We analyse the role played by two different approaches of spatial heterogeneity in theoretical models of annual plants dynamics. The first approach is called quasi-continuous gradient in which one type of resource is changing gradually along the gradient line. In the second one, called the patches approach, part of the habitat is covered by patches and the resource has a different value in each patch. We show that when the spatial heterogeneity of the habitat is small, the two approaches yield the same average number of surviving species, even if a small number of patches is used. In a strong heterogeneity it takes many patches to get similar results as in the gradient case. The difference between the gradient and patchy description of the spatial heterogeneity increases with the number of species present in the system. We have also shown that even when the average number of surviving species is the same, the abundances of species are ordered in a different way, like different species are the dominant ones. The conclusion of this paper is that modelling spatial heterogeneity in a system of plants is not a simple task. Special care is needed when the heterogeneity of the habitat is large, since then depending of the choice of a method, some predictions may differ significantly, making the model non-robust. Therefore the type of theoretical approach must closely match the modelled ecosystem.

## 1 Introduction

Individual-based models provide an important tool to understand the properties of complex ecological and biological systems. To be useful, such models should be robust, i.e. their conclusions should not depend on the particular way some characteristics of the problem are described.

An illustration of this problem is given by modelling of plant dynamics in spatially heterogeneous habitat and the goal of the paper is to compare two commonly used approaches to describe spatial heterogeneity (SH) of the habitat on which several populations of annual plants are growing. Different types of SH are encountered in nature (Stein et al. 2014), but for modelling it two approaches are generally used. In the first one a quasi-continuous gradient in one or more resources (O'Brien 1993; Wilson and Nisbet 1997; Travis et al. 2006) is the source of the SH. In the second one (Tilman 1994; Roxburgh et al. 2004; Silvertown 2004) a number of patches on which the resource has different values, is introduced. The problem of defining heterogeneity is a difficult one, as shown by Stein et al. (2014) and it is important in theoretical and simulation models in view of the recently very active (Kadmon and Allouche 2007; Hortal et al. 2009; Lundholm 2009) topic of the relation between SH and biodiversity (BD). To the best of our knowledge it has never been checked whether the two approaches are equivalent in all or only in some special conditions. Without such a check, statements coming from the results of one type of description have a rather limited validity.

To study this problem we have chosen a version of our model (Droz and Pękalski 2013; Pękalski and Szwabiński 2013) of annual plants and water as the resource.

## 2 Method

The model describes the dynamics of several annual plants competing for a resource (water). We use here an individual-based type model and computer simulation technique. The habitat is a square lattice of linear dimensions *L* × *L* with *L* = 200, having thus 40 000 cells. On each cell there may grow just one plant, or it could be empty. All types of plants have the same demand for water, which will be our stick yard in the sense that the tolerance of plants to the surplus of water and the water falling on each cell (rainfall) will be normalised with respect to the demand, and therefore the tolerance *t*, the rainfall, *w* and the moisture along the gradient, *wg*, as well as the moisture in patches *wp*, will be dimensionless quantities.

The plants are of several types, differing in just one aspect – their tolerance to a surplus of water (Tardieu 2003). To avoid having a system of clones, we allow within each type of plants for small fluctuations of the tolerance. Therefore the tolerance of a plant *i* belonging to the type *m* has the value

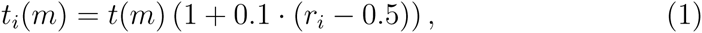

where *r*_*i*_ ∈ (0, 1) here and in the following is a random number taken from a uniform distribution. In the following we shall skip for clarity, the dependence of the tolerance on the type of plants, as in each case it will be evident. The factor 0.1 ensures that we have small fluctuations only, without changes of the type of the plant. In a homogeneous system all cells receive the same amount of water, *w*.

The first type of spatial heterogeneity is introduced as a quasi-continuous gradient of steepness *α* along the *OX* axis when the moisture decreases with increasing *x* coordinate as

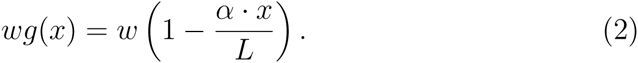

The value of the *y* coordinate is immaterial since all cells for a given value *x* have the same moisture. The gradient is quasi-continuous because the *x* variable changes in a discrete way. With increasing value of *α* the habitat becomes more diversified, hence we assume that the value of *α* is our measure of SH. The total amount of water the system receives, depends on the value of the gradient and is equal to *L*^2^*w*(1 − 0.5 · *α*).

Another type of SH is introduced by putting on the lattice a certain amount, *n*, of square patches of the same size *l* × *l* with different values, attributed randomly, of the moisture on each patch. Outside of the patches the moisture is *w*. The patches are distributed randomly over the lattice. They can touch each other, but cannot overlap. All cells within a given patch have the same amount of water. In order to have correspondence with the case of the gradient, we assume that for a given gradient steepness *α* the amount of water on patches may vary in the same range as for the gradient case, namely between *w* and *w*(1 − *α*). The difference is proportional to the gradient steepness and a cell having e.g. its right top corner at (*x, y*) has the water given by

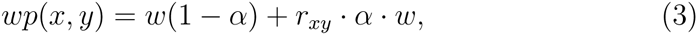

where *r*_*xy*_ is another random number.

Life cycle of the plants is composed of germination and adulthood with seed production phases. Initially we put an equal number (1000) of plants of each species in randomly chosen cells. Plants interact with their nearest neighbours. Namely, the roots of each nearest neighbouring plant (in the von Neumann neighbourhood) block a certain amount of water which could be used by the plant in the center (Schenk 2006). The water available to a plant *i* located at cell (*x, y*) is given by

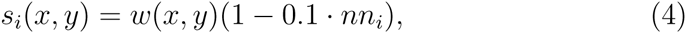

where *nn*_*i*_ is the number of plants in the nearest (von Neumann) neighbourhood and *w*(*x, y*) is the moisture at the cell. Here and in the following *w*(*x, y*) is equal either *wg* or *wp*, depending on the type of heterogeneity considered. The supply, *s*_*i*_ to the tolerance, *t*_*i*_, ratio for the plant *i*

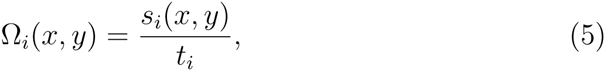

is then used to calculate the probability that the plant *i* survived and can produce seeds

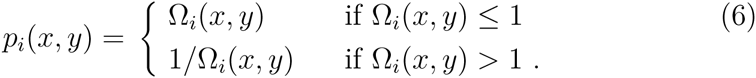

A random number *r*_*i*_ ∈ (0, 1) is chosen and if *p*_*i*_(*x, y*) ≥ *r*_*i*_, the plant passed the survival test and can produce seeds. The number of seeds it produces, *f*_*i*_, depends on *p*_*i*_(*x, y*), as the conditions during the plant’s life determine also its fecundity (Shirley 1929; Seifan et al. 2012)

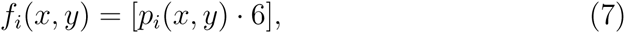

where [(..)] is the integer part of (..) and 6 is the maximum number of seeds a plant can produce in optimal conditions (Seifan et al. 2012). Seeds are then dispersed randomly over 13 cells in the nearest neighbourhood, including the cell on which the plant grows, and then the plant dies and is removed from the system.

Next comes the germination phase. From a cell containing seeds of different species one seed is chosen following the lottery model of Chesson and Warner Chesson and Warner (1981). The fraction of seeds of a given type in the cell determines the probability that a seed of this type will be chosen for germination. Each cell is visited only once. A seed chosen for germination is then put to a germination test with a probability analogous to the one for survival of adults plants, eq.(6), except that there is no blocking of water from the neighbouring seedlings which have too short roots for that. Hence in the germination phase water available to a seed *i* at (*x, y*) is just equal to the moisture at that cell and we define the function 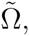, analogous to the one given by eq.(5) as

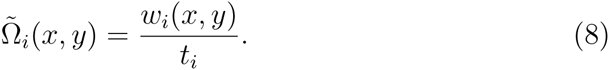

Germination probability, *g*_*i*_(*x, y*) for a seed *i* at the cell located at (*x, y*) has the same form as for the survival of adult plants

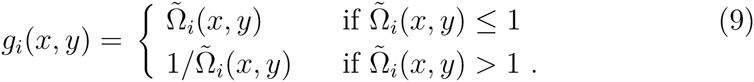

When a seed passed the germination test it becomes a seedling. Once all cells with seeds have been checked, the germination phase is over, the seedlings become adult plants, all seeds are removed from the system (no seed bank) and a new year begins. Plants are randomly chosen, the function Ω is determined. Plants are put to the survival test, produce seeds if they passed the test, disperse them randomly and then die.

The model has the following control parameters:

1. steepness of the gradient *α*,
2. size of the patches *l*,
3. number of the patches *n*,
4. rainfall *w*.

The simulations were carried out for 150 time units (years) when the system reached a stationary state, similarly as in (Seifan et al. 2012) and were averaged over 50 independent realisations, differing by random distribution of initially put plants.

At first we shall study a simple system with just 5 types of plants having tolerances 0.6, 0.8, 1.0, 1.2, 1.4. For such a system we shall demonstrate more detailed features of the total community. Then the problem of equivalence between the gradient and patchy habitats, measured here by the average number of species that survived till the end of simulations, will be studied on a system with 20 and 30 types of species having tolerances differing by 0.1 within the range [0.4,2.3] and [0.4,3.3] respectively.

## 3 Results

In figure 1 we show how the average number of species existing at the end of simulations depends on the value of the rainfall *w*. We take three values of the gradient: *α* = 0.25 (small gradient), *α* = 0.50 (medium) and *α* = 1.0 (large gradient). For a patchy system we shall present two cases (for each value of the gradient): the case of a small number of small patches (10 patches of linear size *l* = 10), called henceforth the case one, and the case of large number of large patches (30 patches of *l* = 20), called case two. We have found that for a sufficiently large patches (*l* ≥ 10) their number is more important than their size.

**Figure 1:**
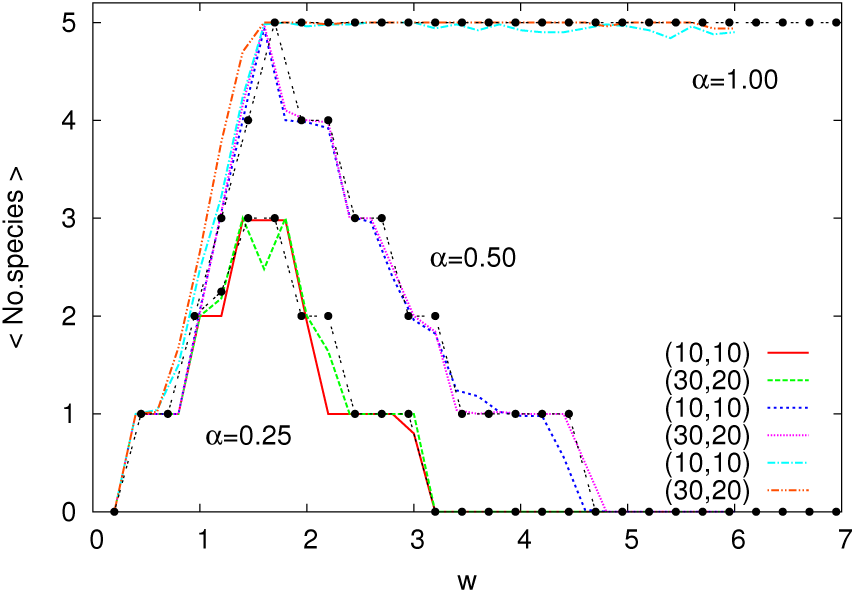
Average number of species surviving till the end of simulations as a function of moisture *w* for 3 values of the gradient steepness *α* and two patchy systems – with small (case one) and large (case two) number of patches. There are 5 species in the system. The data for the respective quasi-continuous gradients are shown with points.

As could be seen from this figure, there is not much difference neither between the small and the large patchy case, nor between the gradient and patchy habitats.

It is interesting to study this system in a more detailed way. Let us start with the time evolution of the 5 species for a given value of the control parameters – *α, l, n, w* and the two cases – case one and case two. The situation for a small gradient is shown in Figure 2.

**Figure 2:**
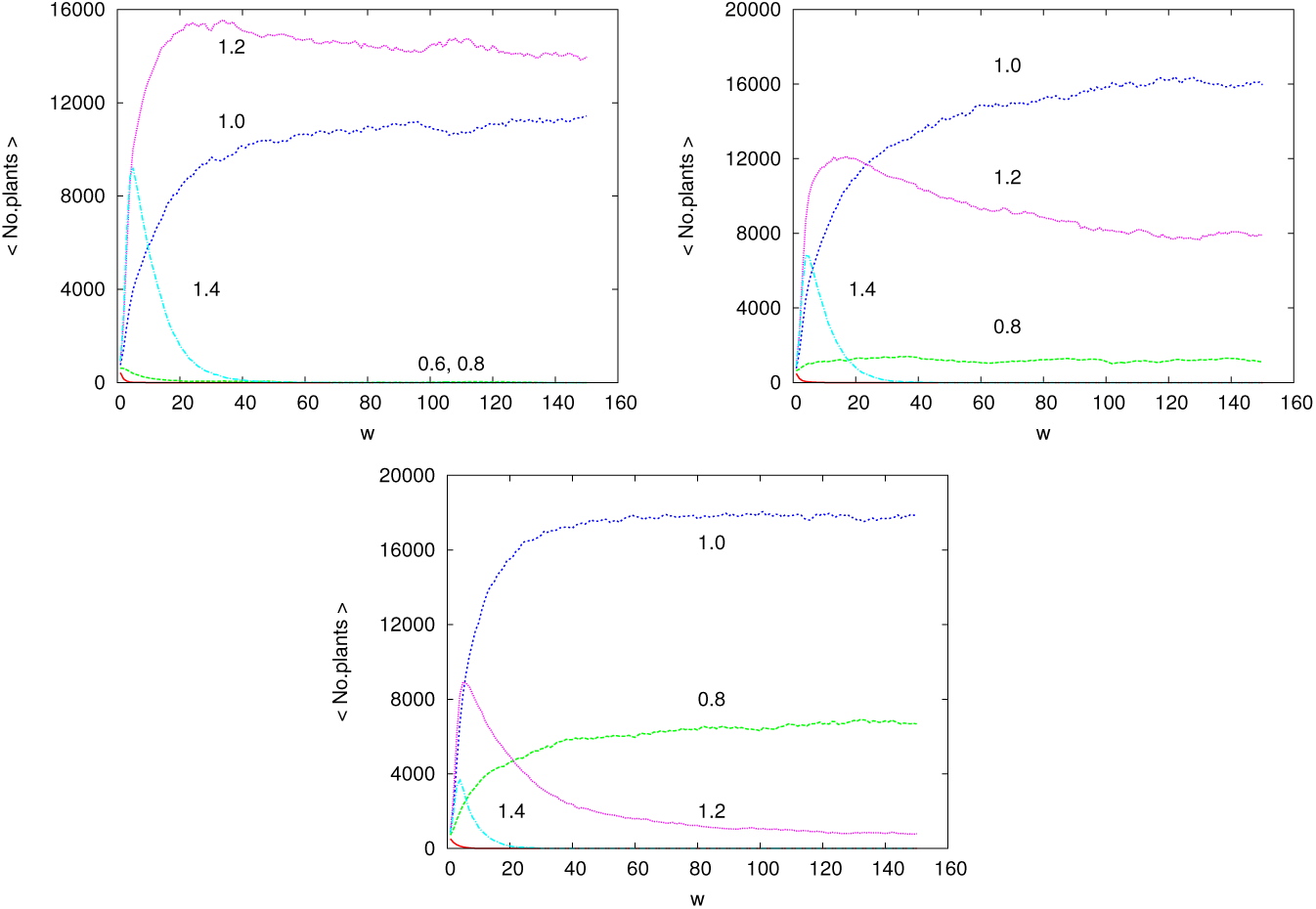
Time evolution of the 5 types of species with tolerances 0.6, 0.8, 1.0, 1.2, 1.4, for *w* = 1.4, for a small gradient *α* = 0.25 and case one (left panel) and case two (right panel). The case of the quasi-continuous gradient is on the bottom panel.

The same dependence but for a medium gradient is shown in Figure 3, while Figure 4 presents the case of the large gradient. In each of the figures, for a given value of *α*, there is the same number of surviving species (4), although for the case of the small number of small patches, the abundances of the weaker species are very low.

**Figure 3:**
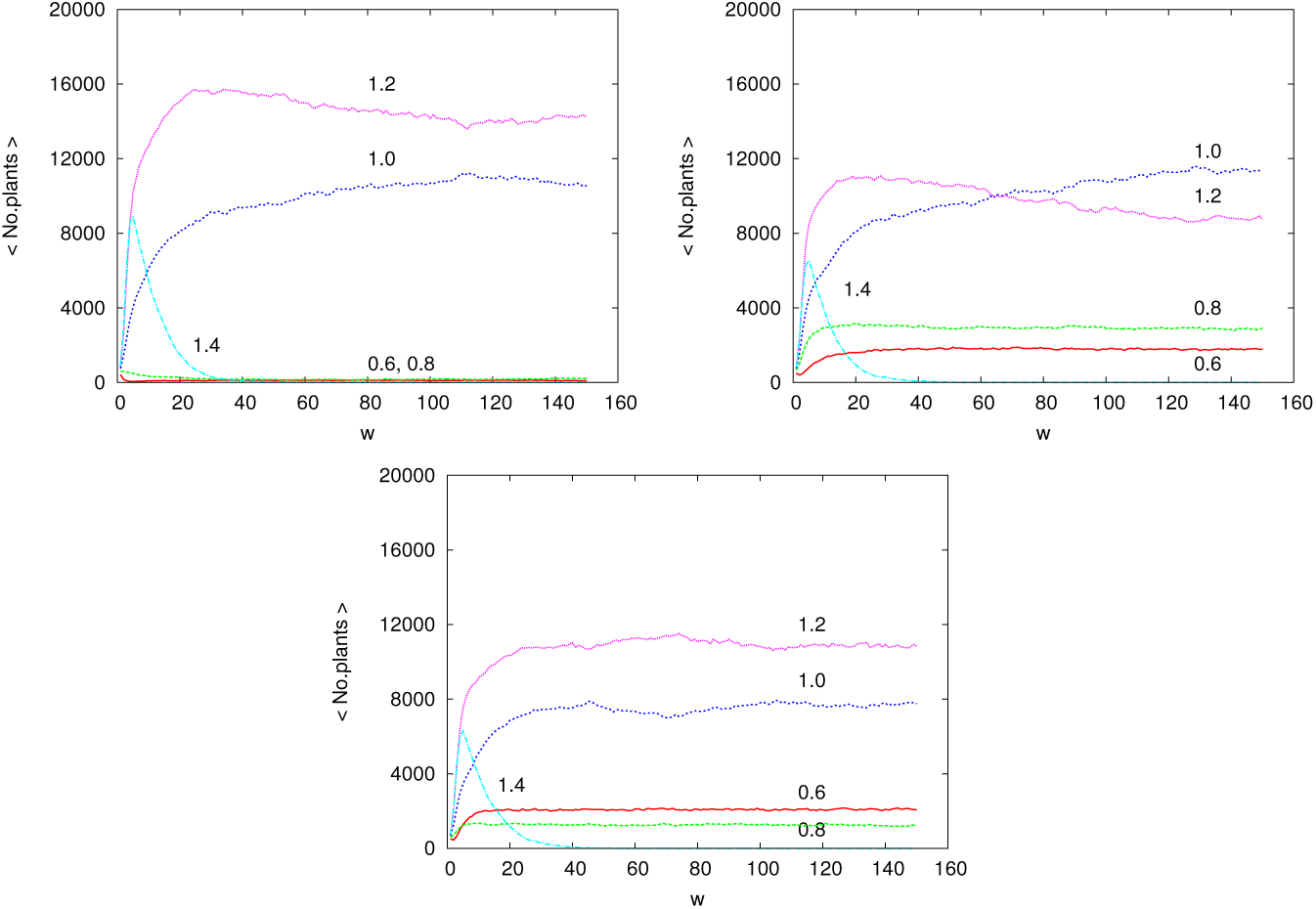
The same as above but for a medium gradient *α* = 0.50.

**Figure 4:**
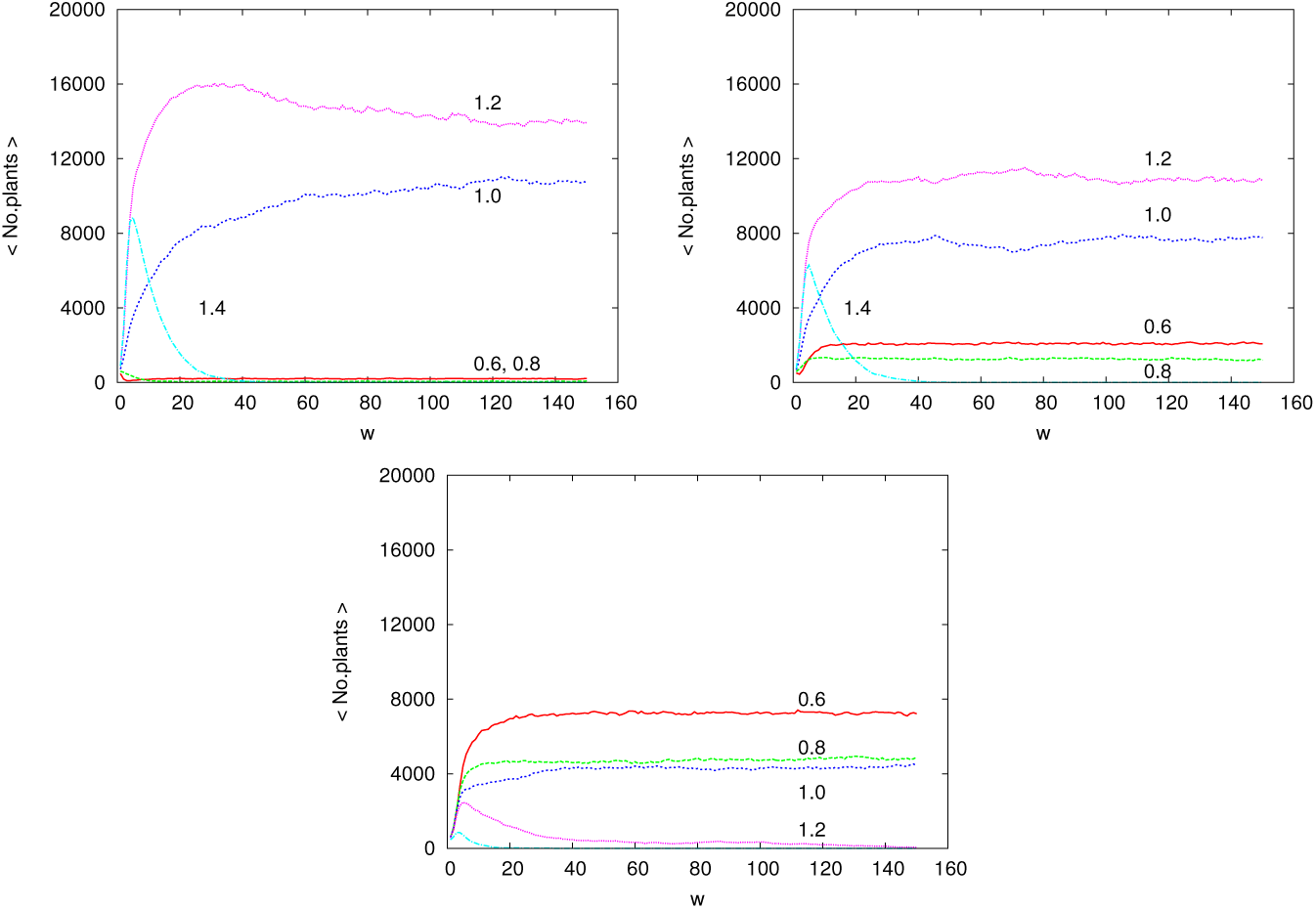
As above, but for the large gradient *α* = 1.0.

As seen from Figure 1 the average amount of surviving species is quite similar for a given value of *α*, for the gradient and two types of patchy systems. However (see Figure 2) the time evolution shows that generally (apart for the plants with highest tolerance which disapear after some time), the dominant species for long time, are not the same for the gradient and the patches cases. The effect becomes even larger when there are more species in the system. It is therefore not justified to claim that the two approaches (gradient and patches) give the same results, even for a very limited number of species.

In Figures 5 we show what are the spatial patterns of the plants at the end of simulations for the case of large gradient (*α* = 1.0) and corresponding large number of large patches. Darker points represent plants with larger tolerance. White areas indicate zones with no plants. We see that when the gradient is quite large, there appear inhabitable zones in both types of description of the SH, and the area suitable for living is reduced.

**Figure 5:**
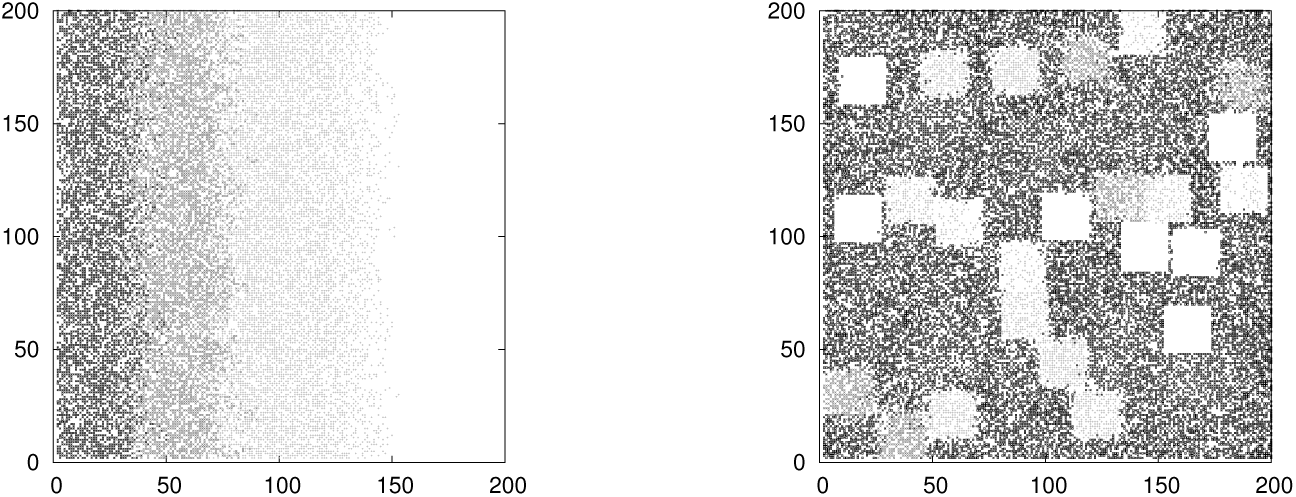
Final distribution of species for *w* = 1.4 and for *α* = 1.0. Left panel shows the gradient, right panel the patches cases.

In order to study more fully the role of the two ways to represent the SH, we need a system with more types of plants. This is done by considering 20 and 30 types of annual plants with tolerances differing by 0.1 and covering the range [0.4,2.3] and [0.4,3.3] respectively. The results are shown in Figures 6 and 7 also for 3 values of *α* and the cases of quasi-continuous gradient and two patchy systems (small number of small patches and large number of large patches).

**Figure 6:**
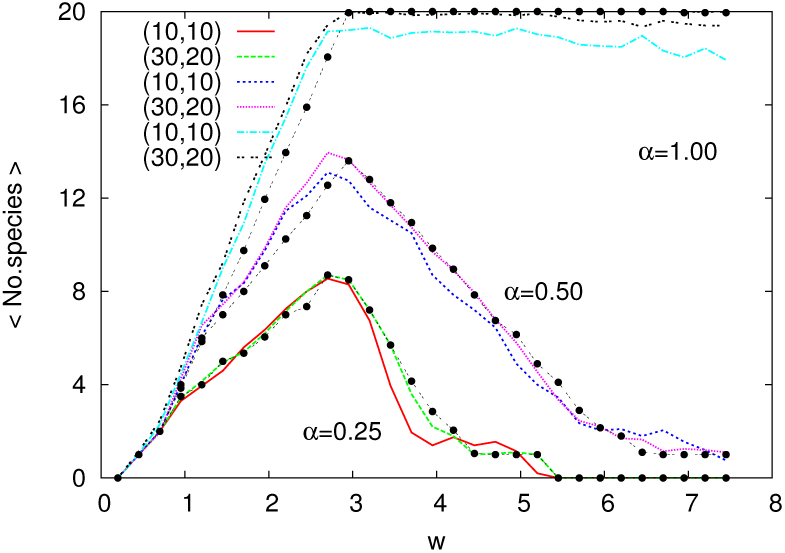
Average number of species surviving till the end of simulations as a function of the moisture *w* for 3 values of the gradient steepness *α* and two systems – case one and case two. There are 20 species in the system. The data for the respective quasi-continuous gradients are shown with points.

**Figure 7:**
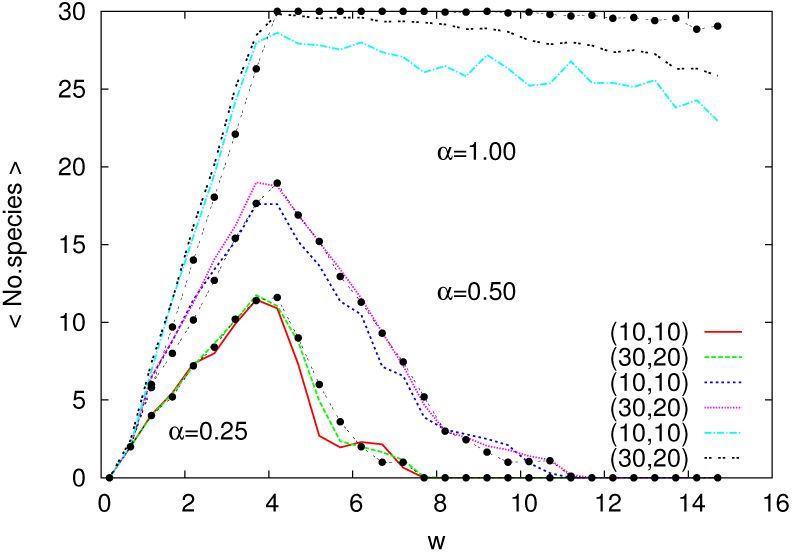
The same as in Figure 6, but for 30 species.

The features of Figures 1, 6 and 7 are quite similar in several aspects – small differences for low values of *α* among the gradient, small and large number of patches cases or increase of the average number of species with increasing *α*. When the gradient is large, the number of species in the system is important. The difference between the gradient and patchy systems, which was negligibly small for 5 species, becomes visible for 20 species and it is still bigger for 30 species. Increasing the number of patches always brings the average number of surviving species in a patchy system closer to those in the gradient one, but as the number of species grows, more and more patches are needed to get similar results.

This similarity between the results coming from a system with relatively small number of species and a one containing 4 or 6 times more of them, indicate that the same type of conclusions would come from much larger systems, hence they have a generic character. Our results show however that different types of plants are dominant when the SH is introduced via the gradient of the resource and different when it is a patchy system.

## 4 Conclusions

We have presented a detailed study of two, commonly used in the literature, ways of introducing SH into a model – via a quasi-continuous gradient of the resource and through a certain amount of square patches with different values of the resource. The amount of water in the gradient case varies between *w* and *w*(1 − *α*) and we assume that in the patchy system water on the patches may vary also in these limits. We have considered three values of the gradient *α* = 0.25, 0.50, 1.0 and systems with small number of small patches and another one with large number of large patches. The value of the gradient steepness *α* is our measure of heterogeneity of the habitat.

We have shown that for small *α* the two approaches (gradient and patches) give quite similar results in the sense that the average number of species surviving till the end of simulations is quite similar and moreover it depends very weakly on the number of used patches. When *α* is large, the agreement between the two approaches depends on the number of species. When there is just a few of them, the agreement, in the average number of surviving species, is reached even when the number of patches is quite small. When there are more types of species, like 20 or more, the results from the gradient and patchy systems are similar only if the number of patches is large. Moreover the number of patches needed to reach the agreement grows with the number of species. We have also demonstrated that the ordering of the species by their abundances is different in the quasi-continuous gradient and patchy habitats, even if the number of species is low and the average numbers of surviving species are the same. This shows that when a more detailed study of the system of plants in heterogeneous condition is performed, the type of description used for SH plays a role and may affect the robustness of the model.

